# Predicting Tissue of Origin from Bulk Tumor Gene Expression using a Pre-trained Transformer Model

**DOI:** 10.1101/2024.12.01.626105

**Authors:** Ted Mellors, Mehran Spitmann

## Abstract

Identifying the tissue of origin for cancers is essential for enhancing diagnostic precision, selecting effective treatments, and guiding clinical decision-making. In this study, we developed a predictive model to classify the tissue of origin across various cancer types. Using a dataset with 10,300 samples from 32 unique tissue types, the model achieved an overall accuracy of 88% in distinguishing among all 32 classes, with an average accuracy of 99.2% within each class. When tested on metastatic skin tumors, it reached an accuracy of 87%, underscoring its potential in addressing challenging metastatic cases. These results demonstrate the model’s reliability in oncology applications, offering a promising tool for improving diagnostic accuracy and supporting personalized cancer treatment strategies.

## Introduction

The accurate identification of the tissue of origin (TOO) for cancers is crucial in oncology, particularly in cases where primary sites are unclear or metastatic presentations occur. This determination informs treatment decisions, facilitates prognostication, and improves overall patient outcomes (Varadhachary et al., 2008; Tothill et al., 2005). Conventional methods for TOO determination often involve clinical evaluations, histopathology, and immunohistochemistry. However, these approaches can be time-consuming, invasive, and, in some cases, inconclusive (Varadhachary et al., 2008).

Advances in high-throughput gene expression profiling, including RNA sequencing, have paved the way for molecular-based TOO prediction. These technologies provide a molecular fingerprint unique to each tissue type, which can be decoded using machine learning algorithms to develop predictive models (Golub et al., 1999; Venet et al., 2011). Machine learning, particularly deep learning, has shown promise in addressing challenges such as the high dimensionality of gene expression data, imbalanced class distributions, and generalization across tumor types (LeCun et al., 2015).

Previous studies have explored various machine learning approaches for TOO prediction, employing feature selection strategies, ensemble methods, and classical algorithms like support vector machines and random forests (Tibshirani et al., 2002; Ramaswamy et al., 2001). However, the need for robust models capable of handling diverse cancer types and metastatic presentations remains unmet.

In this study, we introduce a predictive model leveraging a pre-trained transformer architecture, a state-of-the-art deep learning framework, fine-tuned on gene expression data. Transformers have demonstrated remarkable capabilities in capturing intricate patterns within large datasets, making them well-suited for this application (Vaswani et al., 2017; Theodoris et al., 2023; Mellors, Schneider & Spitmann, 2024). Using data from The Cancer Genome Atlas (TCGA), encompassing 10,300 samples across 32 tissue types, we aimed to develop a reliable tool for TOO prediction that can improve diagnostic precision and inform personalized treatment strategies.

## Results

### Patient Cohort and Data Characteristics

To assess the predictive capabilities of our models, we curated a patient cohort from The Cancer Genome Atlas (TCGA) dataset. Patients were included if they had available gene expression data, primary or metastatic tumor samples, and documented tissue of origin. This resulted in a total of 10,300 patient samples evaluated in this study (**Table 1**).

**Table 1:** Study population and demographics.

| Tissue of Origin |  | Total | Sample Type | AJCC Pathologic Stage |  |  |  |  |  | OS Event | OS |
| --- | --- | --- | --- | --- | --- | --- | --- | --- | --- | --- | --- |
| Abbrev. | Cancer name | n | Primary (Metastatic) | Stage 0 | Stage I | Stage II | Stage III | Stage IV | Unknown | (Yes/No) | Mean (std) |
| ACC | Adrenocortical carcinoma | 79 | 79 (0) |  | 9 | 37 | 16 | 15 | 2 | 28 (51) | 1463.7 (1039.5) |
| BLCA | Bladder urothelial carcinoma | 412 | 412 (0) |  | 4 | 129 | 142 | 135 | 2 | 181 (231) | 796.4 (822.6) |
| BRCA | Breast invasive carcinoma | 1117 | 1110 (7) |  | 184 | 636 | 253 | 20 | 24 | 156 (961) | 1225.2 (1181.9) |
| CESC | Cervical squamous cell carcinoma | 306 | 304 (2) |  |  |  |  |  | 306 | 73 (233) | 1027.3 (1143.6) |
| CHOL | Cholangiocarcinoma | 35 | 35 (0) |  | 18 | 9 | 1 | 7 |  | 18 (17) | 742.5 (548.6) |
| COAD | Colon adenocarcinoma | 480 | 479 (1) |  | 81 | 188 | 133 | 67 | 11 | 104 (376) | 834.5 (749.3) |
| DLBC | Lymphoid neoplasm diffuse large B-cell lymphoma | 48 | 48 (0) |  |  |  |  |  | 48 | 9 (39) | 1346.5 (1469.6) |
| ESCA | Esophageal carcinoma | 185 | 184 (1) |  | 18 | 78 | 56 | 9 | 24 | 78 (107) | 527.1 (481.8) |
| GBM | Glioblastoma multiforme | 157 | 157 (0) |  |  |  |  |  | 157 | 124 (33) | 415.5 (384.8) |
| HNSC | Head and neck squamous cell carcinoma | 522 | 520 (2) |  | 27 | 71 | 81 | 268 | 75 | 221 (301) | 916.4 (868.4) |
| KICH | Kidney chromophobe | 66 | 66 (0) |  | 21 | 25 | 14 | 6 |  | 9 (57) | 2137.6 (1218.2) |
| KIRC | Kidney renal clear cell carcinoma | 541 | 541 (0) |  | 273 | 59 | 123 | 83 | 3 | 175 (366) | 1328.2 (974.9) |
| KIRP | Kidney renal papillary cell carcinoma | 290 | 290 (0) |  | 172 | 21 | 52 | 15 | 30 | 44 (246) | 1054.1 (895.6) |
| LGG | Brain lower grade glioma | 515 | 515 (0) |  |  |  |  |  | 515 | 125 (390) | 962.5 (958.0) |
| LIHC | Liver hepatocellular carcinoma | 371 | 371 (0) |  | 171 | 86 | 85 | 5 | 24 | 130 (241) | 801.0 (727.0) |
| LUAD | Lung adenocarcinoma | 539 | 539 (0) |  | 295 | 126 | 84 | 26 | 8 | 189 (350) | 904.9 (879.3) |
| LUSC | Lung squamous cell carcinoma | 502 | 502 (0) |  | 245 | 162 | 84 | 7 | 4 | 212 (290) | 970.4 (961.4) |
| MESO | Mesothelioma | 87 | 87 (0) |  | 10 | 16 | 45 | 16 |  | 73 (14) | 660.7 (569.7) |
| OV | Ovarian serous cystadenocarcinoma | 421 | 421 (0) |  |  |  |  |  | 421 | 261 (160) | 1182.1 (948.2) |
| PAAD | Pancreatic adenocarcinoma | 179 | 178 (1) |  | 21 | 147 | 3 | 5 | 3 | 93 (86) | 568.1 (476.3) |
| PCPG | Pheochromocytoma and paraganglioma | 181 | 179 (2) |  |  |  |  |  | 181 | 7 (174) | 1057.0 (1155.6) |
| PRAD | Prostate adenocarcinoma | 502 | 501 (1) |  |  |  |  |  | 502 | 10 (492) | 1085.8 (785.3) |
| READ | Rectum adenocarcinoma | 165 | 165 (0) |  | 30 | 51 | 51 | 24 | 9 | 26 (139) | 768.4 (654.9) |
| SARC | Sarcoma | 260 | 259 (1) |  |  |  |  |  | 260 | 99 (161) | 1173.8 (1001.4) |
| SKCM | Skin cutaneous melanoma | 471 | 103 (368) | 7 | 77 | 140 | 171 | 24 | 52 | 222 (249) | 1831.6 (1935.1) |
| STAD | Stomach adenocarcinoma | 412 | 412 (0) |  | 58 | 122 | 169 | 39 | 24 | 158 (254) | 572.5 (530.6) |
| TGCT | Testicular germ cell tumors | 134 | 134 (0) |  | 101 | 12 | 14 |  | 7 | 4 (130) | 2090.8 (1968.3) |
| THCA | Thyroid carcinoma | 513 | 505 (8) |  | 289 | 52 | 113 | 57 | 2 | 16 (497) | 1199.6 (985.0) |
| THYM | Thymoma | 120 | 120 (0) |  |  |  |  |  | 120 | 9 (111) | 1474.2 (1033.5) |
| UCEC | Uterine corpus endometrial carcinoma | 553 | 553 (0) |  |  |  |  |  | 553 | 93 (460) | 1125.7 (897.9) |
| UCS | Uterine carcinosarcoma | 57 | 57 (0) |  |  |  |  |  | 57 | 35 (22) | 871.7 (857.0) |
| UVM | Uveal melanoma | 80 | 80 (0) |  |  | 39 | 36 | 4 | 1 | 23 (57) | 813.6 (551.6) |

### Overall accuracy of tissue of origin prediction

Ensemble models were trained to predict tissue of origin using a robust 10-fold cross-validation approach. To assess model performance across different feature representations, feature sets ranging from 100 to 1000 were tested, alongside a comprehensive model incorporating all available features (“all”). Model evaluation was conducted on both balanced and imbalanced data distributions (**Figure 1.a,b** for balanced data; **Figure 1.c,d** for imbalanced data).

**Figure 1.**
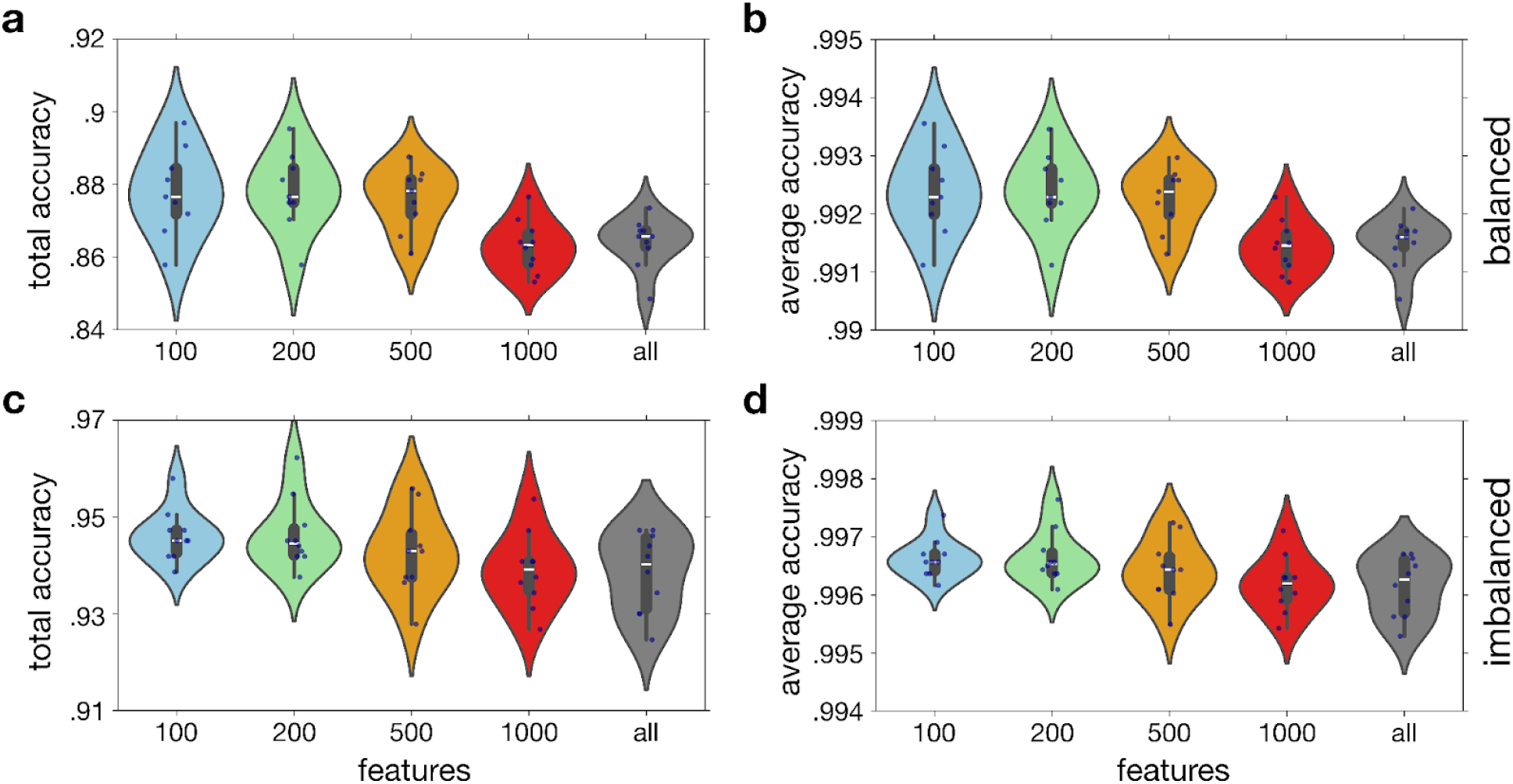
Accuracy of ensemble of classification models with varying numbers of features. Violin plots showing the distribution of total accuracy (**left column**) and average accuracy (**right column**) for balanced (**top row**) and imbalanced (**bottom row**) cross-validation sets. Ensemble models were trained using 10-fold cross-validation. Feature sets ranging from 100 to 1000 were tested, along with a model incorporating all available features (“all”). The central white line represents the median, the box edges represent the interquartile range (IQR), and the whiskers extend to 1.5 times the IQR. Points outside this range are considered outliers.

Two accuracy metrics were used: total accuracy and average accuracy. Total accuracy represents the proportion of correct predictions across all validation samples, while average accuracy measures the mean accuracy achieved across each of the 32 tissue classes.

On balanced data, the model demonstrated a total accuracy of 88% and an average accuracy of 99.2%, highlighting its ability to generalize across tissue types. On imbalanced data, performance improved further, with a total accuracy of 95% and an average accuracy of 99.6%. These results demonstrate the model’s high reliability across diverse tissue types and data distributions, affirming its potential as a robust tool for predicting tissue of origin.

### Accurate Prediction of Tumor Tissue Types Across Diverse Data Distributions

Results illustrate the prediction outcomes for the validation sets in both balanced and imbalanced scenarios (**Figure 2). Figures 2.a** and **Figures 2.b** display results for balanced validation sets, while **Figures 2.c** and **Figures 2.d** show outcomes for the raw, imbalanced test sets.

**Figure 2.**
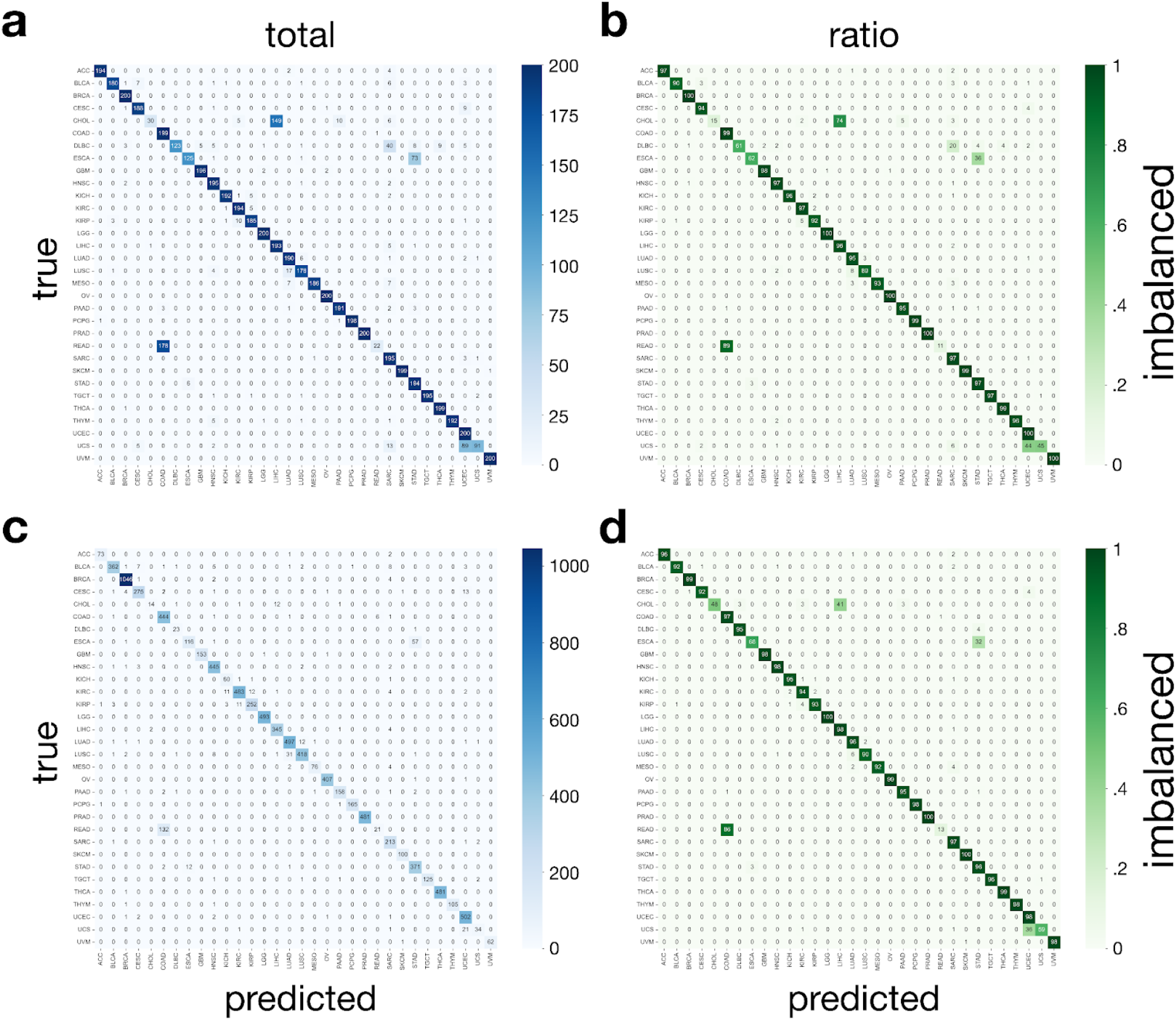
Confusion matrices illustrating classification performance on balanced and imbalanced datasets for selected ensemble models. Total count (**left column**) and ratio (**right column**) of correctly and incorrectly classified instances for balanced (**top row**) and imbalanced (**bottom row**) cross-validation sets. Ensemble models were trained using 100 features with 10-fold cross-validation. Color intensity represents the number of instances (**left column**) or the ratio (percentage values inside the plots, **right column**) for each true class (y-axis) and predicted class (x-axis).

**Figures 2.a** and **Figures 2.c** present the confusion matrices as actual counts of predictions made for each class. These matrices provide detailed insight into the number of times each tissue type was correctly or incorrectly classified, allowing for a granular view of the model’s performance.

**Figures 2.b** and **Figures 2.d**, on the other hand, show the fraction of samples correctly predicted for each class, offering a normalized perspective. This representation helps to understand the relative performance across different classes, regardless of their sample size.

Together, these confusion matrices underscore the model’s ability to handle both balanced and imbalanced datasets effectively, highlighting areas of high accuracy and identifying classes where prediction errors are more common. This comprehensive evaluation reinforces the model’s robustness in predicting tumor tissue types across diverse data distributions.

### Model performance among metastatic test set tumor samples

To assess model accuracy in predicting tissue of origin for metastatic tumor samples, the 100 feature ensemble models were evaluated on a dedicated test set of 351 metastatic tumor samples originating specifically from skin (SKCM) tissue. Due to limited sample size, other metastatic types were excluded from this analysis.

Figure 3. presents detailed prediction results, with panels **Figure 3.a** and **Figure 3.c** showing total counts of predictions for each class, while panels**Figure 3.b** and **Figure 3.s** display the ratio of samples predicted for each class. In balanced evaluation, the model accurately classified SKCM samples 87% of the time. In the raw, imbalanced evaluation, the model achieved an 88% accuracy.

**Figure 3.**
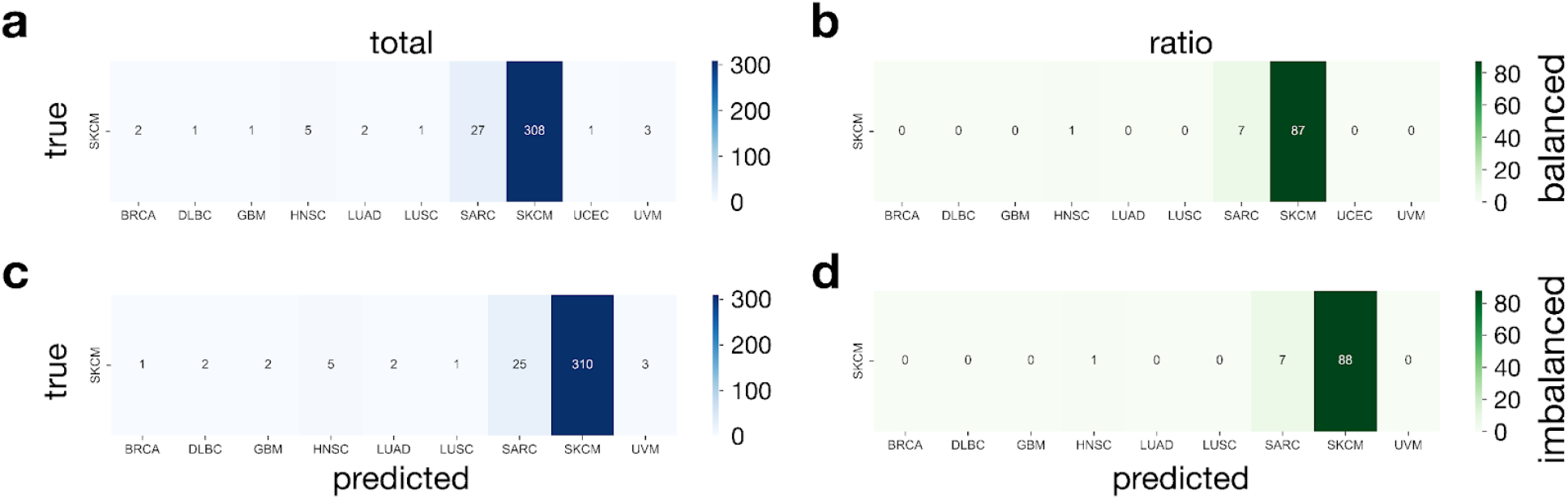
Confusion matrices illustrate the classification performance of a balance and imbalance trained ensemble models on test datasets. Total count (**left column**) and ratio (**right column**) of correctly and incorrectly classified instances for balanced (**top row**) and imbalanced (**bottom row**) models. Ensemble models were trained using 100 features. Color intensity represents the number of instances (**left column**) or the ratio (percentage values inside the plots, **right column**) for true class (y-axis; SKCM) and predicted class (x-axis).

These findings indicate strong model performance in classifying metastatic samples from skin origin, reinforcing its potential for accurate tissue classification in metastatic cases.

The metastatic test set samples were further analyzed to assess whether resection site influenced classification accuracy (**Table 2**). High classification accuracy was consistently maintained across different resection sites, indicating that the model effectively generalizes across diverse metastatic presentations of SKCM. This consistency across varied resection sites underscores the model’s potential in real-world clinical scenarios, where metastatic tumors can originate from multiple sites within the body.

**Table 2:** Model performance stratified by resection site among the 351 metastatic tumors with SKCM tissue of origin.

| Category | Accuracy (%) | n |
| --- | --- | --- |
| Overall | 87.7 | 351 |
| Abdomen | 100.0 | 1 |
| Adrenal gland | 100.0 | 1 |
| Bones of skull and face and associated joints | 100.0 | 1 |
| Brain | 100.0 | 2 |
| Colon | 100.0 | 1 |
| Connective, subcutaneous and other soft tissues of abdomen | 100.0 | 2 |
| Connective, subcutaneous and other soft tissues of head, face, and neck | 75.0 | 4 |
| Connective, subcutaneous and other soft tissues of lower limb and hip | 88.9 | 9 |
| Connective, subcutaneous and other soft tissues of pelvis | 100.0 | 5 |
| Connective, subcutaneous and other soft tissues of thorax | 90.9 | 11 |
| Connective, subcutaneous and other soft tissues of trunk | 70.0 | 10 |
| Connective, subcutaneous and other soft tissues of upper limb and shoulder | 88.9 | 9 |
| Connective, subcutaneous and other soft tissues | 90.9 | 11 |
| Endometrium | 100.0 | 1 |
| Frontal lobe | 100.0 | 2 |
| Jejunum | 100.0 | 1 |
| Liver | 100.0 | 1 |
| Lung | 62.5 | 8 |
| Lymph node | 85.0 | 40 |
| Lymph nodes of axilla or arm | 90.1 | 91 |
| Lymph nodes of head, face and neck | 94.7 | 19 |
| Lymph nodes of inguinal region or leg | 86.7 | 45 |
| Parietal lobe | 100.0 | 1 |
| Parotid gland | 0.0 | 1 |
| Pelvic lymph nodes | 100.0 | 7 |
| Pelvis | 100.0 | 1 |
| Peritoneum | 0.0 | 1 |
| Skin of lower limb and hip | 100.0 | 5 |
| Skin of scalp and neck | 75.0 | 4 |
| Skin of trunk | 90.0 | 20 |
| Skin of upper limb and shoulder | 85.7 | 7 |
| Skin | 93.8 | 16 |
| Small intestine | 87.5 | 8 |
| Spinal cord | 0.0 | 1 |
| Spleen | 100.0 | 1 |
| Thorax | 0.0 | 1 |
| Vagina | 100.0 | 1 |
| Vulva | 100.0 | 1 |

These findings support the model’s robustness in accurately predicting the tissue of origin for metastatic SKCM tumors across heterogeneous resection sites, reinforcing its utility for diagnosing and guiding treatment strategies in complex metastatic cases.

## Discussion

Our study highlights the efficacy of a pre-trained transformer model in accurately predicting the tissue of origin (TOO) for cancers across diverse types. Achieving an overall accuracy of 88% across 32 tissue types, with an average class-specific accuracy of 99.2%, the model demonstrates strong generalizability and robustness. Notably, its 87% accuracy in predicting metastatic skin tumors underscores its potential to address complex clinical scenarios where the primary site is unclear.

The use of pre-trained transformer models in this context is a significant advancement. Originally developed for natural language processing, these models excel at capturing complex relationships and patterns within high-dimensional data. By fine-tuning them on gene expression profiles, we leveraged their ability to discern subtle differences between tissue types, achieving superior performance compared to traditional machine learning methods (Vaswani et al., 2017; Brown et al., 2020; Theodoris et al., 2023; Mellors, Schneider & Spitmann, 2024).

Our findings build upon previous research efforts. Earlier studies have utilized algorithms such as support vector machines, random forests, and ensemble methods for TOO prediction (Ramaswamy et al., 2001; Tibshirani et al., 2002). While these approaches have yielded promising results, they often struggle with issues like imbalanced data distributions and limited generalizability. In contrast, our transformer-based model effectively handles these challenges, as evidenced by its consistent performance across both balanced and imbalanced datasets.

Despite its strengths, the model has limitations. Performance variability across different cancer types and clinical contexts suggests the need for further validation. Testing on independent datasets and conducting prospective clinical studies will be essential to confirm its utility in real-world settings (Kourou et al., 2015). Additionally, the “black box” nature of deep learning models presents interpretability challenges, which could hinder clinical adoption. Future work should focus on integrating explainable AI techniques to enhance model transparency and trust (Doshi-Velez & Kim, 2017).

In conclusion, our pre-trained transformer model represents a significant step forward in TOO prediction. By offering high accuracy, robustness, and generalizability, it has the potential to improve diagnostic precision and guide personalized treatment strategies in oncology. Future research should prioritize clinical validation and the development of interpretability frameworks to maximize the model’s impact on patient care.

## Methods

### Dataset Preparation

Gene expression data for this study were sourced from The Cancer Genome Atlas (TCGA), comprising 10,300 patient samples across 32 unique tissue types. Each sample was normalized and log-transformed to ensure consistency and compatibility with downstream machine learning algorithms. Data were divided into training, validation, and test sets using a stratified split to maintain class balance across splits. An additional metastatic skin tumor dataset, consisting of 351 samples, was curated for external validation to evaluate model performance on metastatic presentations.

### Feature Selection

To optimize model performance, feature selection was performed to identify the most informative gene expression profiles. Features were ranked using variance thresholding, mutual information, and recursive feature elimination (RFE) techniques. RFE was particularly effective in iteratively removing the least important features based on model performance. Selected feature sets ranged from 100 to 1000 genes, alongside a comprehensive set using all available features for comparison. Feature selection was conducted independently within each cross-validation fold to prevent information leakage.

### Model Architecture

A meta-classification framework was implemented, combining three machine learning approaches: neural networks (NN), random forests (RF), and a fine-tuned transformer-based model (Geneformer). Each model was optimized independently to maximize its classification performance:

1. *Neural Networks (NN)*: A feedforward neural network with three fully connected layers, dropout regularization, and ReLU activations. The model was trained using the Adam optimizer with a learning rate of 0.001 and a categorical cross-entropy loss function.
2. *Random Forests (RF)*: An ensemble model consisting of 500 decision trees. Hyperparameters such as the number of trees, maximum depth, and minimum samples per leaf were tuned using a grid search approach.
3. *Geneformer*: A pre-trained transformer model fine-tuned on the TCGA gene expression data (Theodoris et al., 2023). Geneformer’s architecture leverages self-attention mechanisms to capture intricate relationships within high-dimensional gene expression datasets. Fine-tuning involved freezing the initial layers and training the final classification layers using a learning rate of 5e-5.

### Meta-Classifier Construction

Predictions from the NN, RF, and Geneformer models were aggregated using a meta-classifier based on logistic regression. The meta-classifier assigned weights to the predictions of individual models, optimizing overall accuracy by leveraging the complementary strengths of each approach. The meta-classifier was trained on validation set predictions to minimize overfitting.

### Training and Validation

The ensemble framework was evaluated using 10-fold cross-validation. Each fold used 80% of the data for training and 20% for validation. Training procedures ensured that feature selection and hyperparameter tuning were conducted exclusively within each fold to preserve the integrity of validation results.

### Performance Metrics

Model performance was evaluated using the following metrics:

1. *Total Accuracy*: The proportion of correctly classified samples across all tissue types.
2. *Average Accuracy*: The mean accuracy across individual tissue classes, providing a measure of per-class performance.
3. *Confusion Matrices*: Detailed insights into misclassification patterns, visualized as raw counts and normalized fractions.

### External Validation on Metastatic Samples

The final model was tested on an independent set of metastatic skin tumor samples to assess its generalizability to challenging clinical cases. Performance metrics, including total and class-specific accuracy, were computed. Additional analyses evaluated the impact of resection site on classification accuracy to determine the model’s robustness across heterogeneous metastatic presentations.

### Statistical Analysis

Statistical significance of model improvements was assessed using paired t-tests and bootstrapping methods. Confidence intervals for accuracy metrics were computed using 1,000 bootstrap replicates to ensure robust estimates.

### Implementation

All models were implemented in Python using the TensorFlow, Scikit-learn, and Hugging Face libraries. Experiments were conducted on a high-performance computing cluster equipped with NVIDIA A100 GPUs to expedite training and inference. The code and processed datasets are available upon request to ensure reproducibility.

